# Quantifying the role of hyaluronic acid on collagen fiber mechanics via femtosecond laser ablation

**DOI:** 10.64898/2026.10.27.762233

**Authors:** Hannah M. Larson, Faith Miller, Natalie Hawley, Olivia Ward, Alysse DeFoe, Jason A. Burdick, Sarah Calve

## Abstract

Despite therapeutic use of hyaluronic acid (HA) for tendon injuries and tendinopathies, the role of HA in tendon mechanical behavior remains poorly understood. Here, we describe the development of a method that integrates femtosecond laser ablation, second harmonic generation imaging, and mechanical testing to quantify collagen fiber recoil within intact murine tendons under load. Fiber recoil following ablation is used as a measure of collagen fiber sliding and local viscoelastic behavior. Aging and enzymatic depletion of endogenous HA both significantly reduce tendon HA content and decrease elastic, viscoelastic, and total collagen fiber recoil, indicating diminished fiber sliding. To restore HA, low molecular weight methacrylated HA (MeHA) is diffused into tendons and retained through photocrosslinking. MeHA supplementation recovers collagen fiber recoil in HA-depleted and aged tendons, restoring micromechanical behavior toward adult control levels. Quasi-linear viscoelastic modeling further reveals HA-dependent changes in relaxation behavior that are not evident from conventional macroscale mechanical measurements alone. These findings identify femtosecond laser ablation as a powerful platform to quantify fiber-scale mechanics and investigate extracellular matrix contributions to tissue material properties, the results of which specifically isolate HA as an important contributor to tendon collagen fiber sliding and mechanical properties.

## 1. Introduction

Musculoskeletal trauma is a large clinical and economic burden, with half of the 33 million musculoskeletal injuries in the U.S. annually involve tendons or ligaments.^[1],[2]^ Because tendon is primarily comprised of extracellular matrix (ECM) with few cells, healing generally results in incomplete recovery with scar tissue formation and inferior mechanical properties.^[3]^ Tendon injury, pathology, and adverse healing outcomes can cause pain, limit mobility, and decrease quality of life, motivating the need to better understand how ECM composition alters tissue material properties and function.

The ECM provides the mechanical structure necessary for tendon to transmit muscle-generated loads and assists in the cellular communication and biochemical regulation of resident tendon cells.^[4]^ The structure of tendon ECM can be thought of as a fiber-reinforced composite of type I collagen and elastic fibers embedded in an amorphous matrix of proteoglycans (PGs), glycosaminoglycans (GAGs), and glycoproteins.^[5]^ Fibrillar collagens make up 60-85% of tendon dry mass and dominate the mechanical response of tendon across hierarchical levels.^[6]^ However, the non-collagenous ECM that encompasses the other 15-40% of the dry weight also influences the material properties of tendons.^[6]^ PGs and GAGs are hydrophilic, imparting tissue hydration and viscoelasticity, with small leucine rich proteoglycans (SLRPs) and their attached GAGs directly interacting with type I collagen.^[7], [8], [9], [10], [11], [12]^ Hyaluronic acid (HA), a non-sulfated GAG with high binding affinity to water, has been suggested to reside in linear chains between collagen fibrils in ligaments and other connective tissues; ^[13], [14], [15]^ however, the distribution of HA and effect on microscale ECM mechanics of tendons is unclear.

Even though little is known regarding the role of HA in tendon biology and mechanics, the use of HA in the clinical setting is increasing.^[16], [17]^ Injection of HA into joint spaces as a treatment for tendinopathy and tendon injury has been the subject of multiple clinical trials; however, HA is being used more as a biocompatible material rather than a key component of the ECM to modulate tendon physiology.^[18], [19], [20], [21], [22], [23], [24]^ Whether the exogenously applied HA directly penetrated the tissue or impacted local mechanics was not reported in any of these trials. HA used in orthopedic therapies is typically > 450 kDa, and most FDA-approved HA formulations are > 1,000 kDa and/or highly crosslinked.^[19], [25]^ The likelihood of high molecular weight (MW) or crosslinked HA incorporating into tissues is low. Penetration of HA into skin decreased as MW increased, and there was minimal incorporation into the epidermis or dermis for HA > 300–600 kDa.^[26], [27], [28]^ However, as HA is broken down by resident hyaluronidases (HAases) into smaller fragments of 10–100 kDa, the lower MW HA chains are more likely to penetrate into the tissue and affect cellular and microscale mechanics.^[25]^ Knowledge of how HA affects tendon biology, including the local mechanical response, will help elucidate how HA-based therapeutics can facilitate the healing of damaged tendon.

During physiological loading it is thought that microdamage to the ECM can occur, which is repaired by the resident cells to maintain tissue integrity. However, if repair is compromised by pathology or excessive loading, microdamage can accrue and eventually result in tendinopathy.^[29]^ Microdamage and tears to type I collagen fibers have been observed in response to in vivo fatigue in various tendon/ligament model systems.^[30], [31], [32], [33], [34]^ Even though microdamage is not always correlated with immediate changes in macroscale tendon mechanics, it is hypothesized that damage at this scale directly affects cellular behavior and propagates to adjacent cells.^[33], [34], [35]^ The amount and rate of deformation affects the cellular response, and identifying how the composition of the tissue affects the magnitude of ECM deformation around microtears will provide insight into how pathological changes in the ECM, such as those seen in aging^[36]^, lead to tendon injury and disease.^[37]^

At the microscale, continuous interfibrillar shear was observed in tendons, demonstrating load transfer between collagen fibers.^[38], [39]^ It was previously thought that SLRPs surrounding collagen fibrils support this load transfer via noncovalent bonds between sulfated GAG (sGAG) sidechains, but more recent computation modeling implicated friction between collagen fibrils as a mechanism of interfibrillar load transfer.^[7], [40], [41]^ Therefore, it follows that the amorphous ECM that resides in between collagen fibers, which includes HA, may alter the interfibrillar friction through hydration state and fluid viscosity, yet this has not been measured for ECM components in situ.^[41], [42], [43]^ In fatigue injured tendons, HA content increased in the same manner as sGAGs do during tendinopathy, which has been hypothesized to be an adaptation to decrease tissue stress.^[44]^ Enzymatic removal of HA in homeostatic tendons appeared to decrease elastic energy storage and increase the viscous energy dissipation; however, the effects at the fiber scale were not assessed.^[44]^ Furthermore, when hyaluronan synthases *Has1* and *Has3* were genetically knocked out in mice, the maximum stress during pull to failure testing increased in the Achilles tendon.^[45]^ Taken together, the published data led us to hypothesize that HA plays a mechanical role in tendon by modulating interfibrillar friction and shear forces between collagen fibrils, as well as macroscale tissue viscoelasticity.

A specific challenge to measuring the mechanical contribution of HA to tendon material properties lies in a lack of methodology to capture the effect of amorphous ECM components on collagen fiber micromechanical behavior. In this study, we established a method to measure multiscale mechanical behavior in tendon, specifically focusing on collagen fibers and their interactions with surrounding ECM in the larger tendon tissue system. We used two-photon microscopy and femtosecond laser ablation to observe and perturb tendon collagen fibers in their native boundary conditions. By ablating a collagen fiber segment within a tendon under load, we were able to image fiber recoil to resolve differences in microscale material properties.

Our femtosecond laser ablation method was then used to elucidate the mechanical role of HA in tendon (**Scheme 1**). First, we quantified changes in collagen fiber recoil in HAase treated and aged tendons, where the latter are a naturally occurring model of GAG depletion.^[46],[47]^ We hypothesized lower HA content in tendon would reduce bulk tissue stress relaxation and collagen fiber sliding due to decreased tissue hydration, and we found that both groups had decreased HA content that resulted in less collagen fiber recoil. To supplement HA back into tendons, we diffused low MW HA functionalized with methacrylate groups (termed MeHA) and used photocrosslinking to increase the effective HA MW in situ.^[48]^ Finally, we tested whether HA supplementation could restore collagen fiber recoil in HAase-treated and aged tendons. Our results demonstrated that HA supplementation can enhance fiber recoil revealing a previously unknown role for this ECM component in tendon mechanics.

**Scheme 1:**
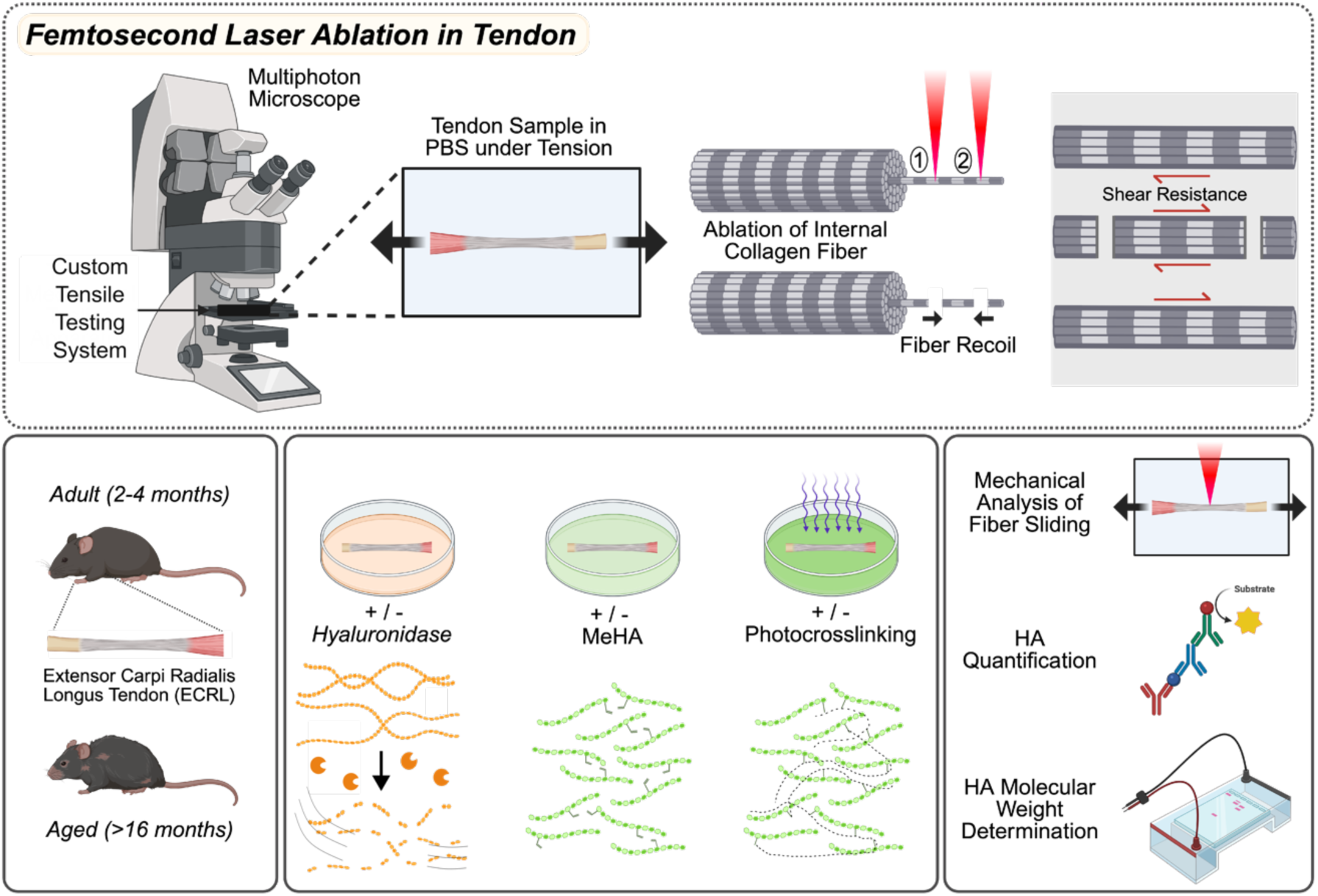
Development of femtosecond laser ablation method to study the role of HA in tendon mechanics. Extensor carpi radialis longus (ECRL) tendons from the murine forelimb were used to test how removal of endogenous HA and supplementation with methacrylated HA (MeHA) affect collagen fiber sliding.

## 2. Results and Discussion

### 2.1. Quantification and Validation of Femtosecond Laser Ablation in Tendon

Extensor carpi radialis longus (ECRL) tendons were harvested from murine forelimbs and specifically selected due to their simple architecture (*i.e.* no higher level organization into fascicles) and their advantageous aspect ratio for tensile loading (∼25:1).^[49]^ Tendons were tested the same day as harvest and tissues at the insertion points (*i.e.* ECRL muscle, metacarpal bone) were preserved. Maintenance of the natural boundary conditions enables testing to occur without directly gripping the tendon tissue for accurate measurement of fiber sliding.

Mechanical testing was performed using our custom 3D printed floating-platform system integrated with an Olympus FVMPE-RS twin laser multiphoton, where tendon samples were submerged in phosphate buffered saline (PBS) and gripped at the muscle and bone insertion sites (**Figure 1A**).^[50]^ The twin laser multiphoton system enables near-simultaneous ablation and imaging, whereas a single laser would require time to tune between wavelengths for each process, preventing capture of the instantaneous mechanical response of the collagen fiber. Fibrillar collagen was visualized using second harmonic generation (SHG) at 910 nm excitation with an Insight X3 laser and ablated at 800 nm excitation with a MaiTai eHP DeepSee femtosecond pulsed laser, taking care to focus below the surface of the tissue. A sequence was performed to 1) displace the tendon at 1% s^-1^ to a set force of 80 mN; 2) hold tendon for 5 minutes of stress relaxation to allow for partial equilibration; 3) take an initial *z*-stack of the tendon; 4) ablate two lines, cutting a collagen fiber segment; and 5) image the recoil of the ablated fiber segment at 0.3 Hz from 100 ms after ablation to 300 s after ablation. A set force of 80 mN was selected as it loaded tendons just past the transition from toe to linear region of the stress-strain curve where all fibers are recruited and load bearing, within the physiological range of motion (2-3% ^[51]^, **Figure S1**). Ablation lines were drawn after stress relaxation using a straight-line region of interest that was the width of a single collagen fiber (∼10 μm). An initial fiber segment was formed by spacing lines 20 μm apart. Image sequences were used to quantify instantaneous (elastic) fiber recoil, time dependent (viscoelastic) fiber recoil, and total recoil (**Figure 1B,C**).

**Figure 1.**
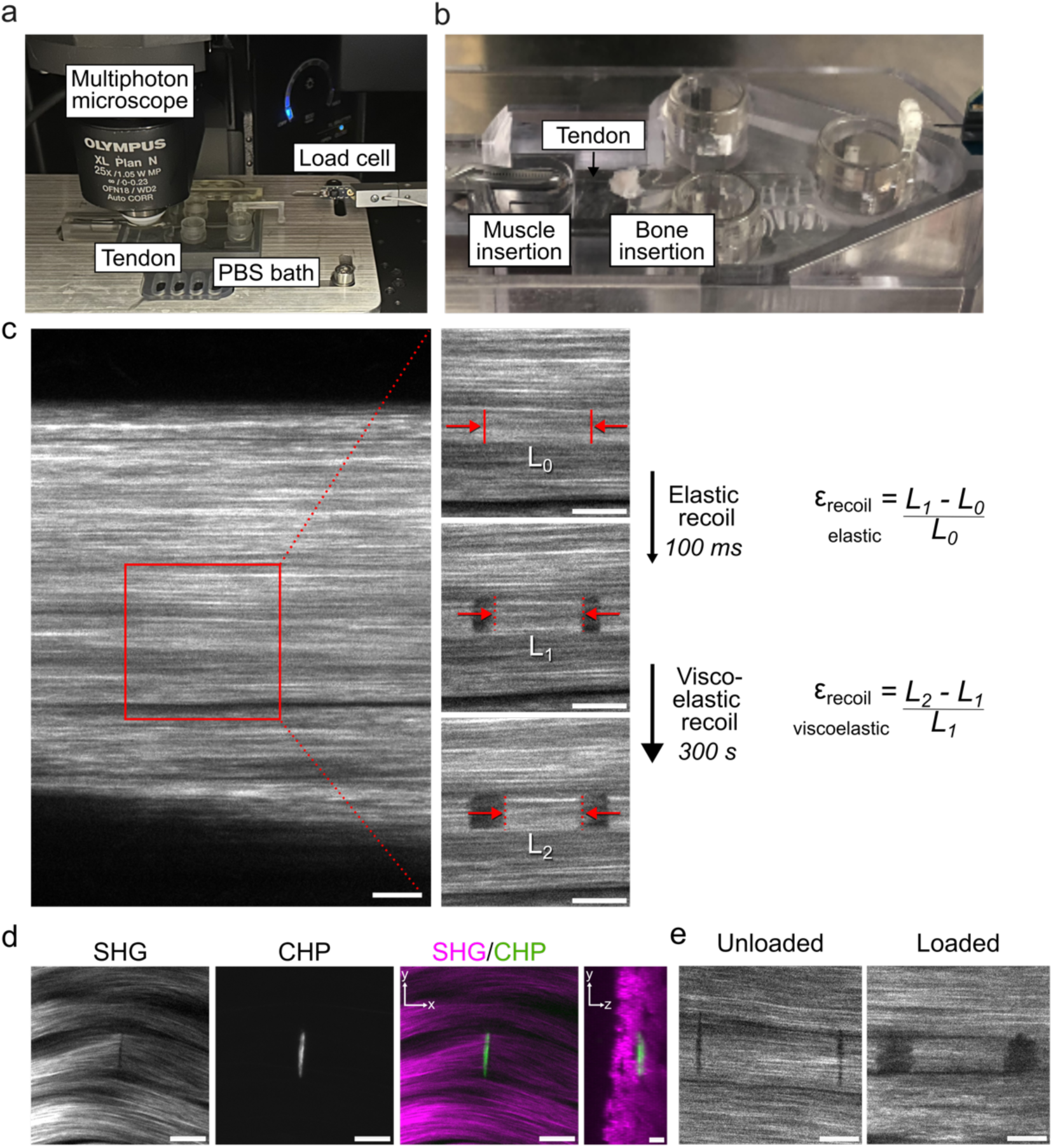
Femtosecond laser ablation selectively severed collagen fibers within tendon and recoil was dependent on mechanical tension. a) Custom 3D-printed testing platform integrated under a multiphoton microscope. b) Close up of tendon and muscle and bone insertions. c) SHG image of tendon under tension; insets show individual collagen fiber where ablation occurred (solid red lines, L_0_), elastic fiber recoil (dashed lines, L_1_), and viscoelastic fiber recoil (dashed lines, L_2_). d) Ablation was indicated by lack of SHG signal, and verified using CHP staining, with orthogonal projection showing CHP^+^ cut is beneath the surface of the tendon. e) Ablation in unloaded or loaded tendon. All images representative of *n* = 3 biological replicates. All scale bars = 10 µm.

After executing an ablation line, SHG signal no longer appeared where the line was drawn, indicating successful ablation occurred due to the need for organized fibrillar structure to induce SHG (**Figure 1D**).^[52]^ The extent of damage around an ablation site evaluated using fluorophore-conjugated collagen hybridizing peptide (CHP), which selectively binds to denatured collagen.^[53]^ Collagen damage aligned closely with the ablation line. An orthogonal view of the tendon z-stack confirmed ablation selectively damaged the chosen collagen fiber, about 20 μm beneath the tendon surface, but not surrounding tissue. The pulse duration used for laser ablation influences the mechanism by which molecular bonds are broken. Femtosecond pulses, common to multiphoton microscopy and used in this study, initiate a plasma phase change that is well suited for precise cutting, avoiding mechanical side effects of longer pulse durations such as cavitation bubbles and high temperatures that denature surrounding proteins.^[54]^ While nanosecond ablation has been used to create a hole in tendon as a model of tissue microdamage to study tendon healing,^[55]^ our study is the first instance using femtosecond laser ablation to characterize the micromechanical properties of tendon, with ablation features an order of magnitude smaller.

To validate that fiber recoil was dependent on tension, ablation of fiber segments was compared between slack tendons and those loaded to the set force. No fiber recoil after 5 min was observed in unloaded tendons, whereas there was consistent, time-dependent recoil in loaded tendons (**Figure 1E**). These results were in line with previous studies that leveraged pulsed laser energy to characterize mechanical forces during embryogenesis and homeostasis, where recoil magnitude and direction were correlated with the force held in a structure after laser ablation.^[52], [56], [57], [58], [59]^

### 2.2. Effect of Aging and Endogenous HA Removal on Fiber Recoil

To validate that our method could quantify differences in tissues with different ECM content, we used aged tendons, a model in which GAGs are known to be naturally dysregulated. During aging, sGAGs and water content decrease,^[60], [61], [62]^ which occurs concomitant with an increase in tissue stiffness and decrease in viscoelasticity.^[63], [64], [65], [66], [67]^ These reported changes lead us to hypothesize that HA content influences the alterations in mechanical properties observed during aging.

Adult and aged tendons were imaged with biotinylated hyaluronic acid binding protein (HABP) and DAPI to visualize the distribution of HA and cell nuclei, respectively (**Figure 2A**). HABP staining in adult tendons demonstrated a more specific morphology running between arrays of tenocytes and greater signal intensity than aged tendon. To confirm the specificity of HABP staining, we removed endogenous HA in adult tendon using HAase from *Streptomyces hyalurolyticus* to measure whether this disruption would replicate mechanical changes seen with aging. *S. hyalurolyticus* HAase is specific for HA and does not act on sGAGs.^[68]^ HABP imaging of wholemount tendons treated with HAase showed little fluorescence after digestion (**Figure 2B**). Differences in HA content as a function of age and HAase treatment were quantified using an HA ELISA. HAase digested adult tendons had significantly reduced HA compared to adult (**p<0.003, Tukey’s post-hoc) (**Figure 2C**), confirming that HAase effectively reduced native HA content. In addition, aged tendon had significantly less HA per tissue weight than adult tendon (*p=0.02, Tukey’s post-hoc) (**Figure 2C**).

**Figure 2.**
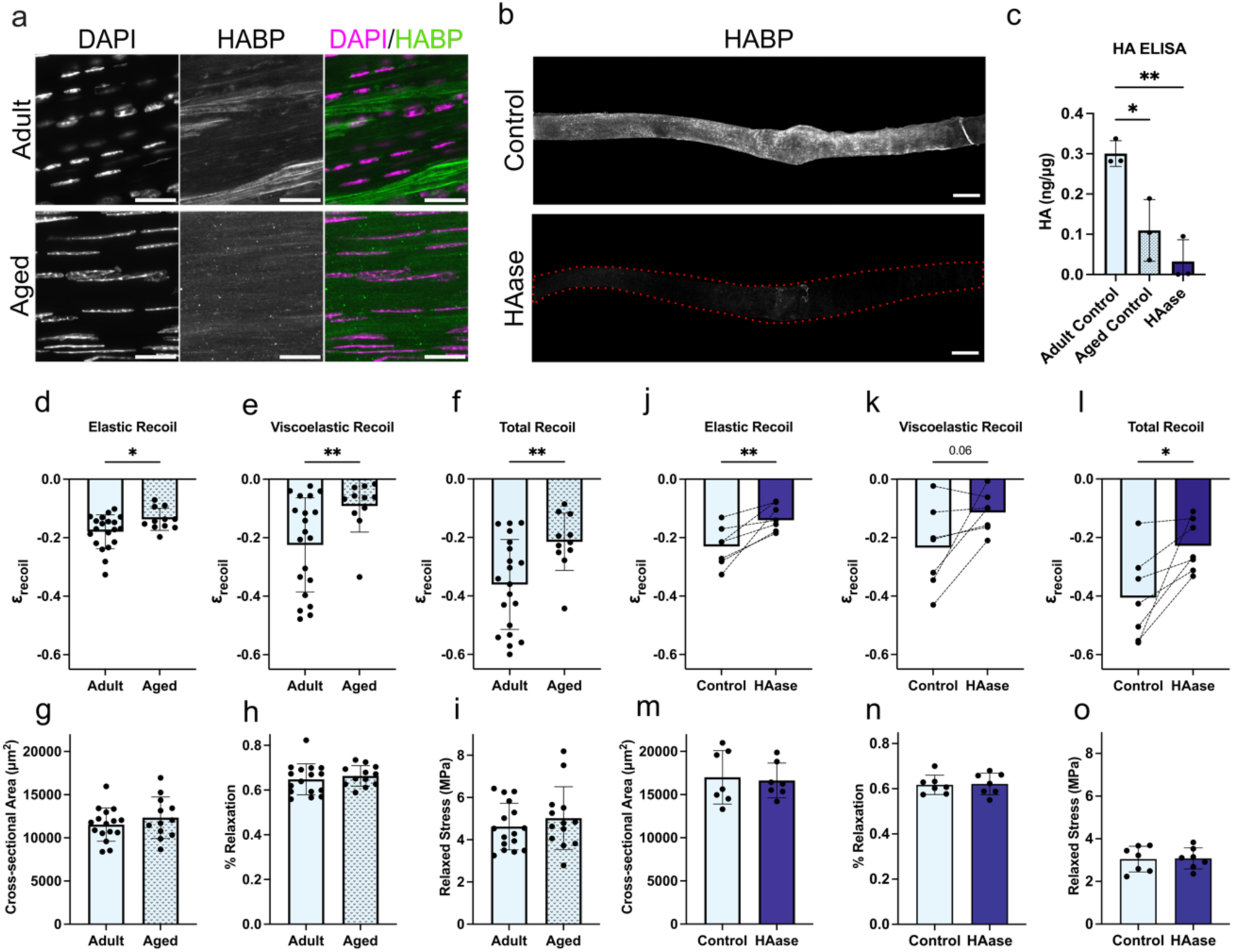
Reduced HA content in aged and HAase-treated tendons reduced collagen fiber recoil. a) Cryosection fluorescent images of HABP and DAPI in adult and aged tendon. Scale bars = 25 µm. All images representative of *n* = 3 biological replicates. b) Wholemount images showing HABP^+^ staining in control and HAase treated tendon. Scale bars = 200 µm. All images representative of *n* = 3 biological replicates. c) HA content per tendon dry weight of adult, aged, and HAase tendons (**p<0.01, One-way ANOVA, Tukey post-hoc test), *n* = 3 per group. d) Elastic recoil of ablated collagen fibers in adult and aged tendon. e) Viscoelastic recoil of ablated collagen fibers in adult and aged tendon. f) Total recoil of ablated collagen fibers in adult and aged tendon. g) Tendon cross-sectional area at ablation site. h) Percent of initial stress after 300 s stress relaxation. i) Stress after relaxation at time of ablation. (d-i) Data represent mean ± SD. Welch t-test, *n* = 21 adult, *n* = 11 aged. j) Elastic recoil of ablated collagen fibers in adult control and HAase treated tendons. k) Viscoelastic recoil in adult control and HAase treated tendons. l) Total recoil in adult control and HAase treated tendons. (j-l) Data represent mean with dashed lines connecting contralateral tendons from single biological replicate. Paired t-test, *n =* 7 per group. m-o) Macroscale measurements of cross-sectional area, percent stress relaxation, and relaxed stress during ablation. Data represent mean ± SD. Welch t-test, *n* = 7 per group. *p<0.05, **p<0.01, ***p<0.001.

Using our femtosecond laser ablation method, adult (3-5 mo.) and aged (>16 mo.) murine ECRL tendons were mechanically tested to assess differences in fiber- and macro-scale mechanical properties. Elastic recoil of collagen fibers was significantly decreased in aged tendons (*p<0.05, Welch’s t-test) (**Figure 2D**). Similarly, viscoelastic recoil was significantly decreased in aged tendon (**p<0.01, Welch’s t-test) as was total recoil (***p<0.001, Welch’s t-test) (**Figure 2E,F**). The reduction in fiber recoil with aging suggests that there is reduced fiber sliding in aged tendon compared to adult, which may play a role in previously published studies showing stiffer and less viscoelastic macroscopic mechanical properties.^[63], [64], [65], [66], [67]^

In contrast, no differences in macroscopic measurements were observed. Cross-sectional area (CSA), actuator displacement from preload to set force, percent stress relaxation, and relaxed force during ablation were the same for both adult and aged tendon (**Figure 2G-I**). Though this result did not match expectations that aged ECRL tendons would be stiffer than in the adult, it is possible that differences may be resolved at higher stresses, whereas our testing set force was lower to resolve microscale differences.

Mechanical testing of HAase digested samples revealed similar differences to those observed for aged tendon. Elastic recoil was significantly decreased in HAase digested tendons compared to contralateral controls (**p<0.01, pairwise t-test) (**Figure 2J**). Similarly, viscoelastic recoil was decreased in HAase digested tendons (p=0.06, pairwise t-test) as was total recoil (*p<0.05, pairwise t-test) (**Figure 2K,L**). These results present a previously unknown role of HA in tendon material properties, suggesting that the depletion of HA increases interfibrillar shear forces and decreases fiber sliding. Like aged tendons, no differences were found in CSA, actuator distance to set force, percent relaxation, or relaxed force during ablation when HA was removed (**Figure 2M-O**). The observation that there were no significant differences in the macroscopic properties highlights the importance of studying multiscale mechanics. These results show that although HA did not have a resolvable effect on macroscale tensile mechanics of the ECRL tendon when loaded to 80 mN, it may still impact tenocyte behavior through the modulation of shear forces at the microscale. However, the ECRL is a positional tendon, and the results may differ for an energy storing tendon such as the plantaris. Furthermore, HA was shown to be enriched in the interfascicular matrix of the Achilles tendon, a more complex tendon with higher order structure,^[69]^ and we expect that the macroscale mechanics will be affected by HAase treatment at this scale. Nevertheless, these data support our hypothesis that HA content influences tendon microscale mechanics and reveals HA content decreases with aging.

### 2.3. Reintroduction of HA into Tendon Tissue

We then investigated how to successfully reintroduce HA into tendon tissue to test if we could recover the collagen fiber recoil in the HAase and aged tendons towards that of the control adult. The relationship between HA MW and penetration into tendon tissue was assessed using fluorescently tagged HA of varying MW (50-1000 kDa). Low MW HA (50 kDa) had the highest degree of penetration, as indicated by the greatest fluorescence, and fluorescent intensity decreased with increasing MW (**Figure 3A**). Although high MW HA (>1,000 kDa) is most commonly used in therapeutics, our results indicate these therapeutics likely do not enter tendon tissue or need to be degraded into smaller fragments prior to diffusion. Despite these results, HA therapeutic trials in tendon have generally reported positive results including reduced tendon gliding resistance, swelling, and inflammation.^[17], [19], [70], [71]^ We hypothesize that these benefits come from interactions with the tendon sheath and epitenon or are achieved after HA chains are broken into lower MW fragments.^[72]^

**Figure 3.**
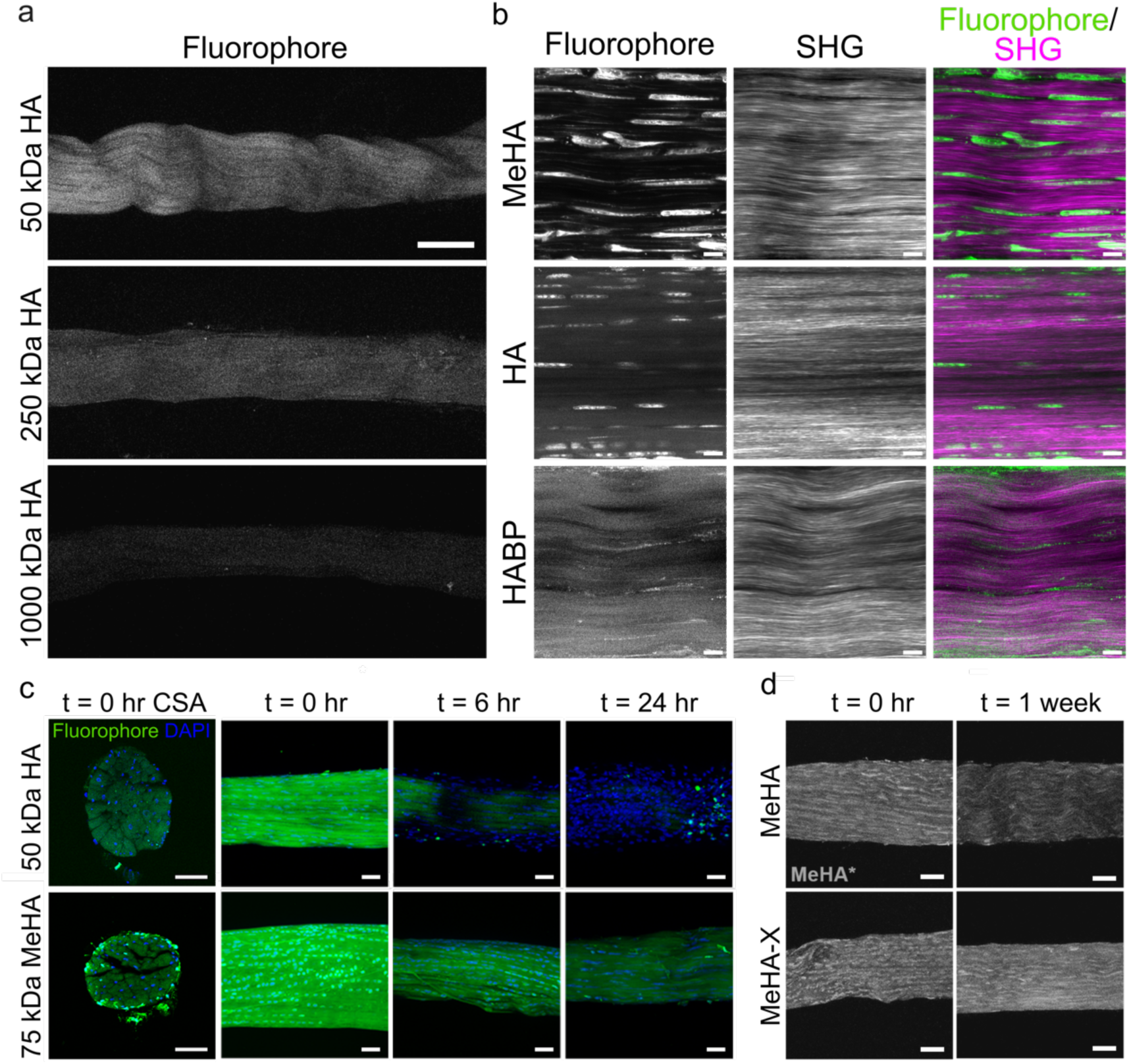
HA diffusion into tendon was best achieved with low MW HA and MeHA, where persistence time increased via methacrylation and crosslinking. a) Diffusion of 1% fluorescently tagged HA of varying MW into tendon. Scale bar = 100 µm. b) Morphology of exogenous MeHA and HA diffused into tendon tissue compared with endogenous HA labelled by HABP in context of collagen SHG. Scale bar = 10 µm. c) Cross-section of tendon after MeHA or HA incubation (top). Whole-mount fluorescence of MeHA and HA residence in tissue at t = 0, 6, and 24 hours after incubation while tendons were hydrated in PBS (bottom). Scale bars = 50 µm. d) MeHA fluorescence in tendon without (MeHA) or with crosslinking (MeHA-X) at t = 0 h and t = 1 week. Scale bar = 50 µm. All images representative of *n* = 3 biological replicates.

To further investigate the effect of HA MW on fibril mechanics, we used methacrylated HA (MeHA) as an alternative for HA supplementation with the goal of diffusing low MW HA into tissue and then effectively increasing the HA MW through crosslinking.^[48]^ The introduction of a methacrylate group to the HA backbone enables MeHA macromers to react with each other via a photoinitiated reaction. MeHA has been frequently used to produce hydrogels with tunable mechanical, viscoelastic, and degradation properties by leveraging HA MW, degree of modification, and concentration.^[48], [73]^ In addition, MeHA was previously shown to diffuse into cartilage to modify cartilage material properties through photocrosslinking.^[74], [75]^

We used two-photon microscopy to validate that the distribution of HA and MeHA diffused into tendon was similar to that of endogenous HA stained with HABP. MeHA was localized in linear arrays as well as around cell bodies (**Figure 3D**). HA staining was similar, though mostly found around cell bodies. HABP staining had notably lower signal than the fluorescently tagged MeHA and HA, but the morphology also appeared enriched around cell bodies in between collagen fibers. The similarities in distribution seen via imaging suggested that supplemented HA would have similar mechanical effects as endogenous HA.

Then, we compared the penetration of 70 kDa MeHA with 50 kDa HA by imaging cryosections of the tendon cross-section, and both were able to diffuse through the full thickness of the ECRL within 1 hour (**Figure 3C**). HA- and MeHA-incubated tendons were then imaged at 0, 6, and 24 hours after incubation to measure the HA persistence in hydrated tissue. Fluorescent 50 kDa HA diffused out of the ECRL quickly, with little fluorescence remaining after 6 hours. Uncrosslinked 70 kDa MeHA dissipated out of the tissue more slowly, with reduced fluorescence 6 hours after treatment and further decreased fluorescence at 24 hours. The fact that MeHA, without crosslinking, stayed in tendon tissue longer than HA suggests that the attached methacrylate group interacts with endogenous ECM, which warrants further study.

The effect of photocrosslinking on MeHA persistence was investigated by incubating ECRL tendons in MeHA (0.5 wt%, 20% methacrylation) and photoinitiator (0.05 wt/vol% lithium phenyl-2,4,6-trimethylbenzoylphosphinate, LAP) for one hour, then crosslinking with 450 nm laser light exposure for 3 minutes. Samples without and with light exposure were imaged after incubation (t=0 h) and again after 1 week of storage in PBS at 4 °C. At 1 week, crosslinked MeHA (MeHA-X) persisted over un-crosslinked MeHA (MeHA), as indicated by higher fluorescence (**Figure 3D**). Thus, the combination of MeHA and light exposure could be a useful therapeutic tool to modulate HA residence time within tissues. Numerous variables can be used to alter features of MeHA diffusion, persistence and degradation, such as the extent of modification of the HA backbone, HA molecular weight, and MeHA concentration.

### 2.4. Supplementation of Tendons with MeHA

#### 2.4.1. Mechanical Characterization of MeHA in Tendon

To test if the observed mechanical differences in HAase digested samples could be reversed, HA was reintroduced by incubating tendons in MeHA for 1 hour. In addition, we investigated the effects of prior HA content and crosslinking (**Table 1**). We employed low concentrations of MeHA (0.5 wt%, 70 kDa) with a low degree of methacrylation (6% modification) to limit crosslinking as endogenous HA is found as linear chains. Control tendons either received no treatment or the photoinitiator plus light exposure, which were not significantly different (**Figure S2)**.

**Table 1:** Sample Groups and Treatments for Mechanical Testing.

| Group | Treatment 1<br>(1 h at 37 °C) | Treatment 2<br>(1 h at RT) | Treatment 3 |
| --- | --- | --- | --- |
| Adult Control | Buffer-only | PBS | No treatment, or LAP + 3 min laser |
| MeHA with Crosslinking (MeHA-X) | Buffer-only | MeHA | LAP + 3 min laser exposure |
| MeHA w/o Crosslinking (MeHA) | Buffer-only | MeHA | No treatment |
| HAase | Buffer + 10 U HAase | PBS | No treatment |
| HAase + MeHA-X | Buffer + 10 U HAase | MeHA | LAP + 3 min laser exposure |
| HAase + MeHA | Buffer + 10 U HAase | MeHA | No treatment |

Mechanical differences between treatment groups were examined as described above, and contralateral tendons that underwent different treatments were compared pairwise. Differences in HA content with treatment were also assessed using an HA ELISA (**Figure S3**). MeHA supplementation resulted in significantly greater elastic recoil than controls (*p<0.05, pairwise t-test), and HAase + MeHA also significantly increased elastic and total recoil (*p<0.05, pairwise t-test), indicating that 70 kDa MeHA increased fiber sliding. Unexpectedly, adult tendons that only underwent MeHA-X treatment showed no difference compared to controls, whereas those digested with HAase and then supplemented with MeHA-X had significantly increased elastic and total recoil compared to samples digested with just HAase (*p<0.05, pairwise t-test) (**Figure 4B,D**). We hypothesize that crosslinking MeHA may form a robust network that negates the fiber sliding effects of uncrosslinked HA; however, tensile testing of MeHA-X tendons using 70 kDa MeHA with 6, 20, and 100% methacrylation did not yield significantly different material properties (**Figures S1, S4, S5)**. These results demonstrate that MeHA supplementation, both with and without crosslinking, after HAase treatment, effectively restored elastic and total fiber recoil to control values.

**Figure 4.**
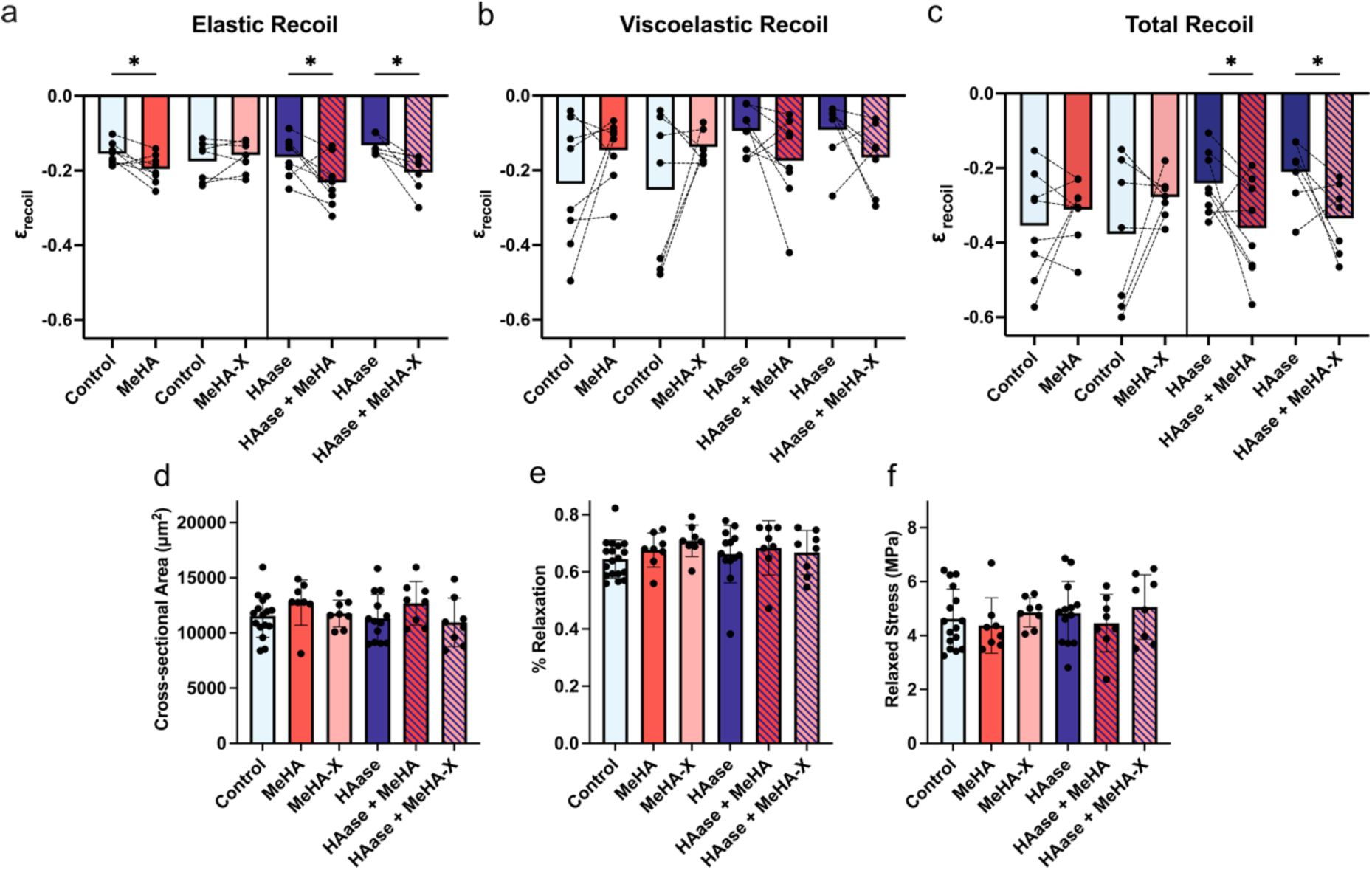
MeHA supplementation recovered fiber sliding behavior in HA-depleted tendons. a-c) Elastic, viscoelastic and total recoil of ablated collagen fibers. Data represent mean with dashed lines connecting contralateral tendons from a single biological replicate. *Brown-Forsythe one-way ANOVA*, *Fisher’s LSD pairwise post-hoc*, *n =* 6-8 per group. d-f) Macroscale measurements of cross-sectional area, actuator displacement, percent stress relaxation, and relaxed stress at ablation. Data represent mean ± SD. *Brown-Forsythe One-way ANOVA, Dunnett’s T3 post hoc, n =* 6-8 per group. *p<0.05, **p<0.01, ***p<0.001.

Viscoelastic recoil did not differ significantly between any tendon treatment (**Figure 4C**). This result did not match expectations that amount and MW of HA would alter the viscoelasticity of the tissue.^[76]^ The variability seen in viscoelastic recoil may be due to the radial position of the ablated fiber within the tendon, as well as other ECM and cells besides what was captured by SHG. An additional contribution to variability may be due to the buffer solution. While the elastic response of buffer-incubated tendons was similar to control tendons not incubated in buffer or treated with LAP or laser exposure (**Figure S2A**), the buffer incubation may have altered tissue hydration or composition.

Macroscale measurements of CSA and actuator displacement to the set force were not significantly different between test groups (**Figure 4E,F**). Similarly, the percent stress relaxation after 300 s and the relaxed stress at the time of ablation were not significantly different with any treatment, which did not support our initial hypothesis that HA supplementation would increase tissue viscoelasticity (**Figure 4G,H**). This result may be attributed to differences in viscoelastic properties that may not have been resolved with our approach. Previous work has also shown that percent relaxation, quantified by only comparing endpoint measurements, does not completely capture viscoelastic behavior, and looking at relaxation rates or other parameters can reveal differences in viscoelastic properties.^[63]^ This dataset was also analyzed for sex differences (**Figure S6**). Elastic recoil was significant for sex/treatment interaction (*p<0.05, two-way ANOVA), and cross-sectional area was significantly different between male and female mice (**p<0.01, two-way ANOVA). These results match expectations that male mice have larger tendons than female, albeit without significantly different material properties.^[77]^

#### 2.4.2. QLV Modeling of HAase and MeHA

To further examine time-dependent changes in viscoelasticity with HAase and MeHA treatment, macroscale stress relaxation and measurements of fiber recoil length were each fit to a Fung Quasi-Linear Viscoelastic (QLV) model, which captures both an elastic component and viscoelastic component of the material response.^[78]^ Parameters of the model, *C_1_* and τ*_1_*, are reported and represent the extent of relaxation (relaxation ratio) and the relaxation time constant, respectively, over the 300 s relaxation window for each response. An additional relaxation ratio and time constant (*C_2_* and τ*_2_*) were calculated to fit the longer time window of macroscale stress relaxation, though only the first terms are reported here. Average QLV parameters for each group were plotted to compare relaxation profiles (**Figure 5A**). Compared to control tendons, MeHA-X and HAase groups had a significantly lower relaxation ratio, *C_1_,* at the macroscale (*p<0.05, ***p<0.001, Fisher’s LSD test; **Figure 5B**). In contrast, there were no differences in relaxation time constant, *τ_1_*, between any groups (**Figure 5C**). These results indicate that stress relaxation occurs over the same time frame; however, HAase or MeHA-X treatments reduce the extent of that relaxation. We hypothesize that the reduction in MeHA-X stress relaxation resulted from the effects of crosslinking, while reduced stress relaxation from HA removal was caused by decreased water content and tissue viscosity.^[62], [79]^

**Figure 5.**
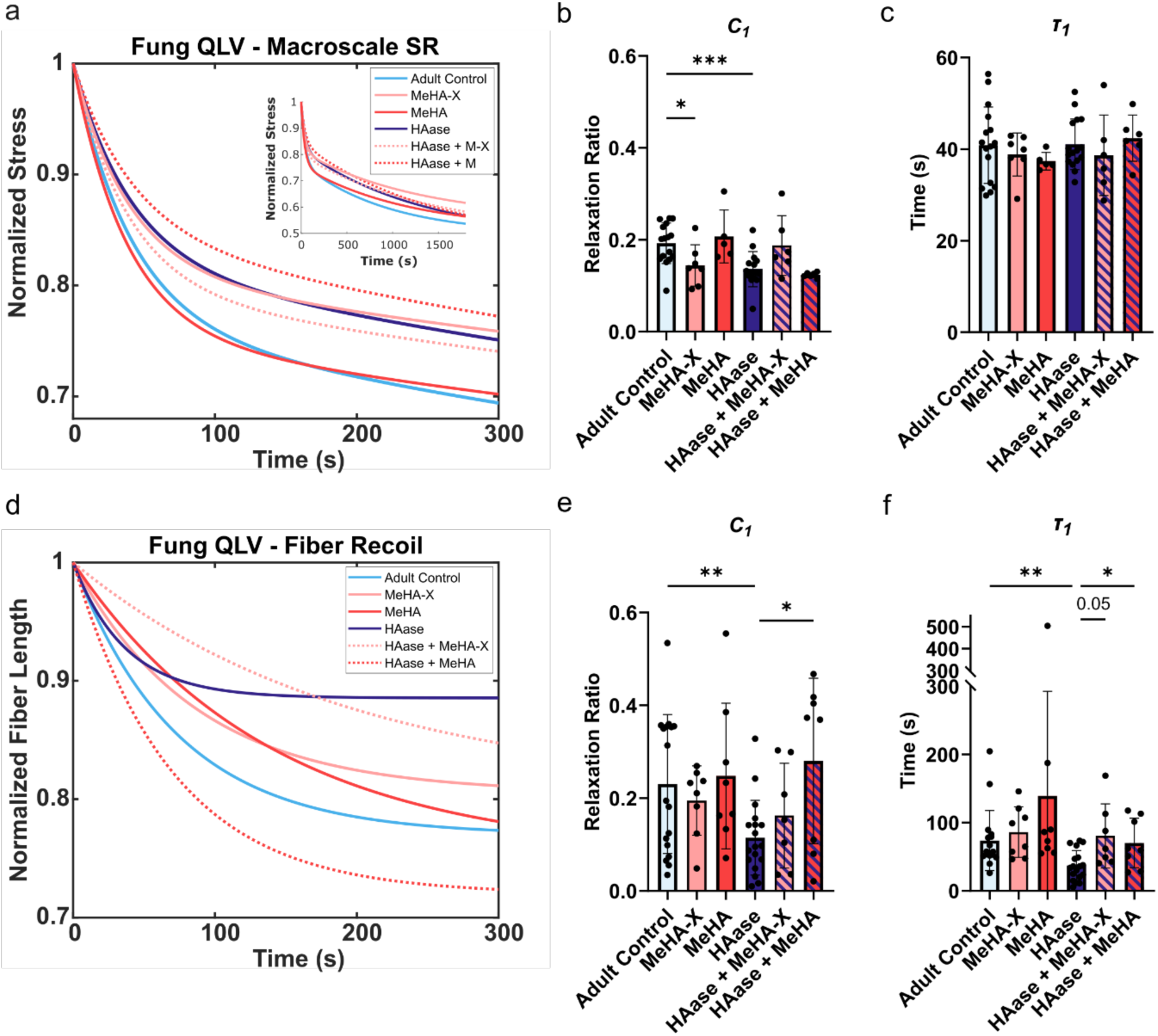
QLV modeling reveals time dependent differences in viscoelasticity of HAase and MeHA-X treated tendons. a) Average QLV fit for macroscale stress relaxation for each group. b-c) Macroscale relaxation ratio and time constant for short-term stress relaxation. Data represent mean ± SD. *Brown Forsythe one-way ANOVA*, *Fisher’s LSD post-hoc*, *n* = 5-16 per group. d) Average QLV fit for fiber scale recoil after ablation for each group. e-f) Fiber scale relaxation ratio and time constant. Data represent mean ± SD. *Brown Forsythe One-way ANOVA*, *Fisher’s LSD post-hoc, n* = 8-18 per group. *p<0.05, **p<0.01, ***p<0.001.

Next, the average QLV fit for fiber retraction of each group was plotted for comparison (**Figure 5D**). At the fiber scale, HAase digestion significantly reduced the relaxation ratio and time constant compared to controls, while MeHA-X and MeHA supplementation did not (**p<0.01, Fisher’s LSD test) (**Figure 5E,F**). Addition of MeHA to HAase digested tendons significantly increased *C_1_* and *τ_1_* (*p<0.05, Fisher’s LSD test), comparable to control levels. In contrast, MeHA supplementation alone did not change macroscale or fiber-scale parameters from controls, and HAase digestion altered macroscopic relaxation behavior that was not recoverable by MeHA supplementation. HA in native tissues is often found complexed with other PGs such as lecticans,^[80]^ which were likely removed during HAase digestion. It could also indicate distinct effects of HA MW on macroscale viscoelasticity, which may be due to the MW of native HA being much larger than the 70 kDa MeHA. Taken together, QLV modeling at the macro and microscale revealed effects of HAase and MeHA treatment on tendon multiscale viscoelasticity that was not captured through the end-point measurements of percent stress relaxation or viscoelastic fiber recoil (**Figure 4A**).

### 2.5. Supplementation of Aged Tendons with MeHA

#### 2.5.1. Mechanical Characterization of MeHA Supplemented Aged Tendons

Aged tendons had a similar decrease in HA content and fiber recoil compared to control adult tendons treated with HAase digestion. Therefore, we next investigated whether MeHA supplementation of aged tendons would restore the fiber sliding behavior to that of control adult tendons. Aged tendons were treated with either MeHA or MeHA-X to compare to adult macroscale measurements and fiber recoil. MeHA significantly increased elastic, viscoelastic, and total fiber recoil compared to aged controls, returning recoil to levels that of adult tendon (*p<0.05, **p<0.01, pairwise t-test). Conversely, MeHA-X treatment significantly decreased elastic recoil and total recoil (**p<0.01, *p<0.05, pairwise t-test) and approached significance for decreased viscoelastic recoil (p=0.05, pairwise t-test) (**Figure 6A-C**). These results differed from MeHA-X in adult tissue and suggest that photocrosslinking impacts aged tissue differently than in adult. The differences between MeHA and MeHA-X in adult and aged tissue may be a result of different composition in tendons with age. Aged tissue may have increased binding interactions with MeHA than adult tissue, whether from increased AGE crosslinks, altered PG and sGAG amounts, or other structural and compositional changes.^[81]^ Macroscopically, MeHA treatment resulted in significantly higher percent relaxation than control tendons (*p<0.05, Fisher’s LSD test), while CSA and relaxed stress at time of ablation were not significantly different between groups (**Figure 6**).

**Figure 6.**
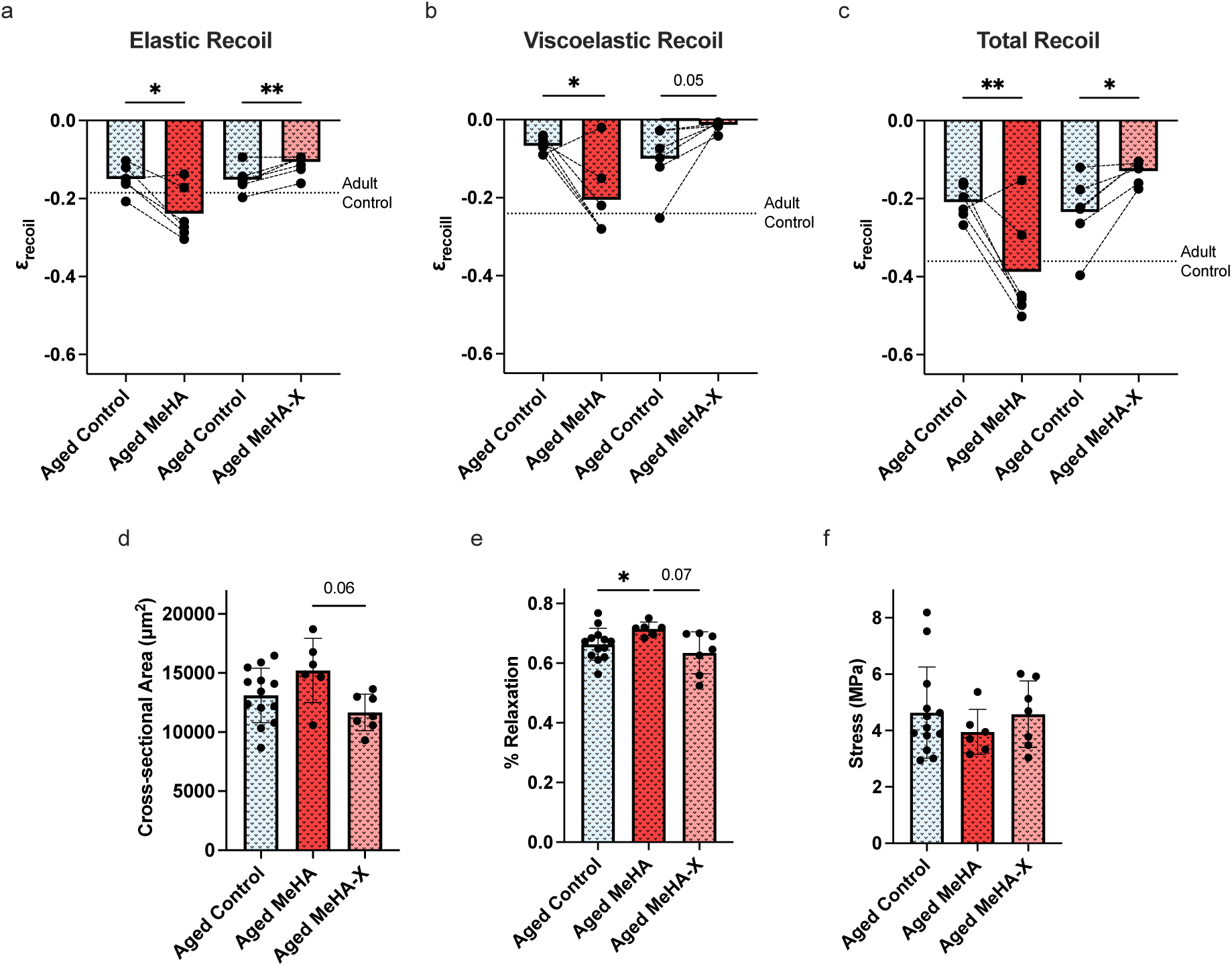
MeHA treatment of aged tendons restored fiber sliding to adult levels. a-c) Elastic, viscoelastic, and total recoil of ablated collagen fibers. Data represent mean with dashed lines connecting contralateral tendons from a single biological replicate. *Brown-Forsythe one-way ANOVA, Fisher’s LSD pairwise post-hoc*, *n =* 6 per group. d-f) Macroscale measurements of cross-sectional area, actuator displacement, percent stress relaxation, and relaxed stress during ablation. Data represent mean ± SD. *Brown-Forsythe one-way ANOVA, Dunnett’s T3 post hoc, n =* 6 per group. *p<0.05, **p<0.01, ***p<0.001.

#### 2.5.2. QLV Modeling of Aged Tendon with MeHA

As done with adult tendons, the viscoelastic response of aged tendon groups was fit to a QLV model to further investigate time-dependent changes. At the macroscale, aged tendon had slightly less stress relaxation than control adult tendon, though not as reduced as HAase digested adult tendons (**Figure 7A**). Contrary to expectation, aged tendons that received MeHA treatment had a significantly reduced relaxation ratio compared to aged controls (***p<0.001, Fisher’s LSD test), and reduced relaxation time constant as well (**Figure 7B,C**). At the fiber scale, aged control and MeHA-X groups had reduced fiber recoil similar to that of adult HAase digested tendons (**Figure 7D**). MeHA-X treatment of aged tendon significantly decreased the relaxation ratio (*p<0.05, Fisher’s LSD test), while MeHA significantly increased the relaxation ratio to adult control levels (**p<0.01, Fisher’s LSD test) (**Figure 7E**). The relaxation time constant for all aged groups was, on average, lower than that of adult controls, though not significantly different between groups. Overall, QLV parameter differences in aged tendon groups matched the differences seen in end-point measurements. Taken together, these results suggest that some mechanical changes in aged tendon may be due to decreased HA or sGAG content and can be recovered with HA supplementation.

**Figure 7.**
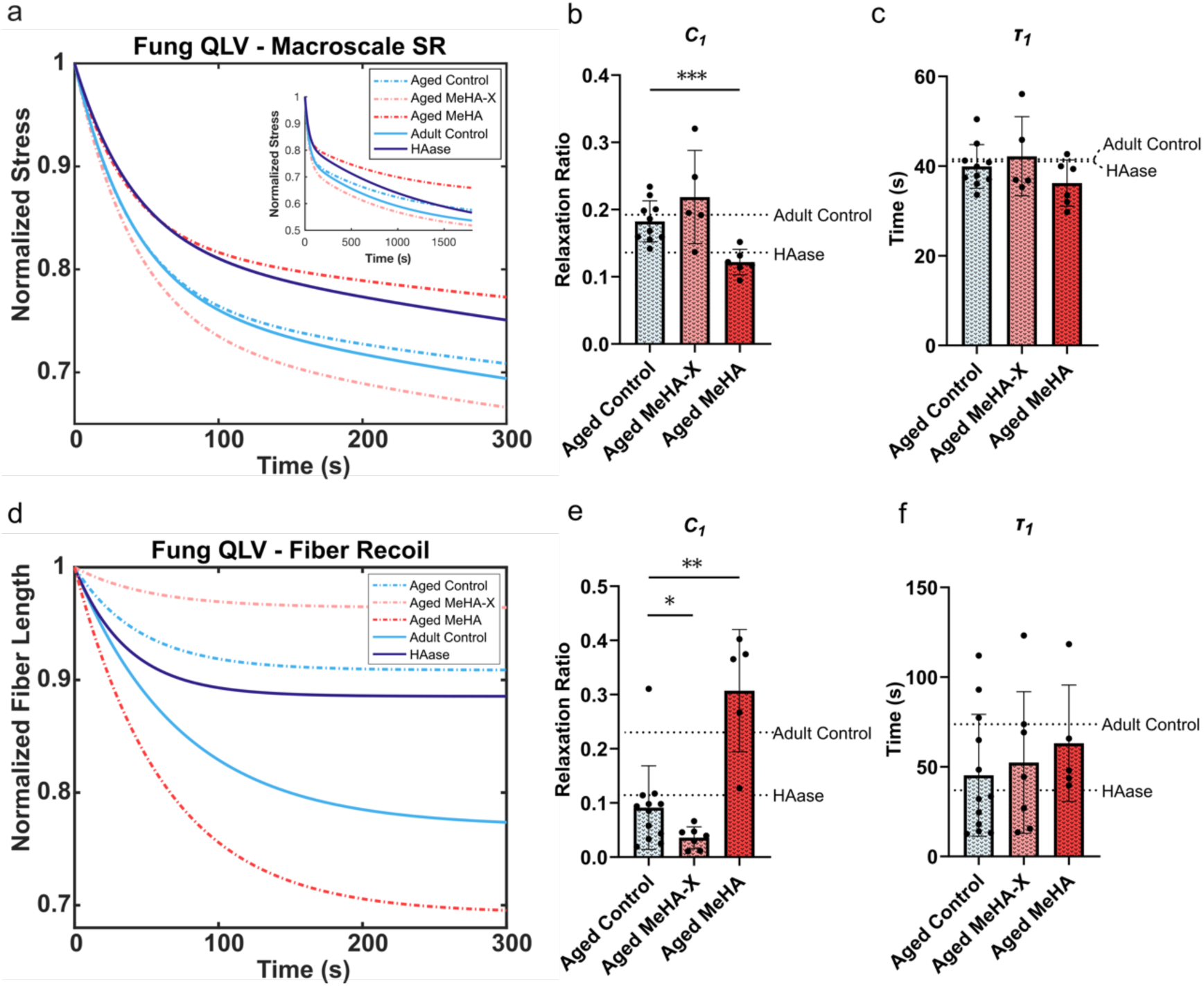
QLV parameters for aged tendon groups matched changes seen in endpoint viscoelastic measurements. a) Average QLV fit for macroscale stress relaxation for each group. b-c) Relaxation ratio and time constant for short-term stress relaxation. Data represent mean ± SD. *Brown Forsythe one-way ANOVA*, *Fisher’s LSD post-hoc*, *n* = 5-10 per group. d) Average QLV fit for fiber scale recoil after ablation for each group. e-f) Fiber scale relaxation ratio and time constant. Data represent mean ± SD. *Brown Forsythe one-way ANOVA*, *Fisher’s LSD post-hoc, n* = 4-9 per group. *p<0.05, **p<0.01, ***p<0.001.

### 2.6. MeHA Crosslinking Study

To better understand mechanical differences brought about by different concentrations of MeHA in the tendon, we altered the time of incubation. Increasing incubation time from 1 hour to 24 hours yielded more MeHA in the tissue, as indicated by higher fluorescent intensity at 24 hours, and similar distributions of MeHA with both incubation times (**Figure 8A**). Samples incubated for 1 hour and 24 hours were photocrosslinked and mechanically tested to ascertain how the dose of MeHA altered crosslinking and mechanical properties (**Figure 8B-D**). Elastic recoil was significantly reduced after 24-hour incubation compared to 1-hour (*p<0.05, pairwise t-test); however, there were no significant differences for viscoelastic and total recoil. These results suggest a more highly crosslinked network forming with longer incubation, leading to stiffer material properties. We expect that increasing the degree of modification of MeHA will yield similar results.^[82]^ From this, we hypothesize that the differential results of MeHA-X treatment in adult and aged tissue may be attributed to MeHA forming additional crosslinks in aged tissue.

**Figure 8.**
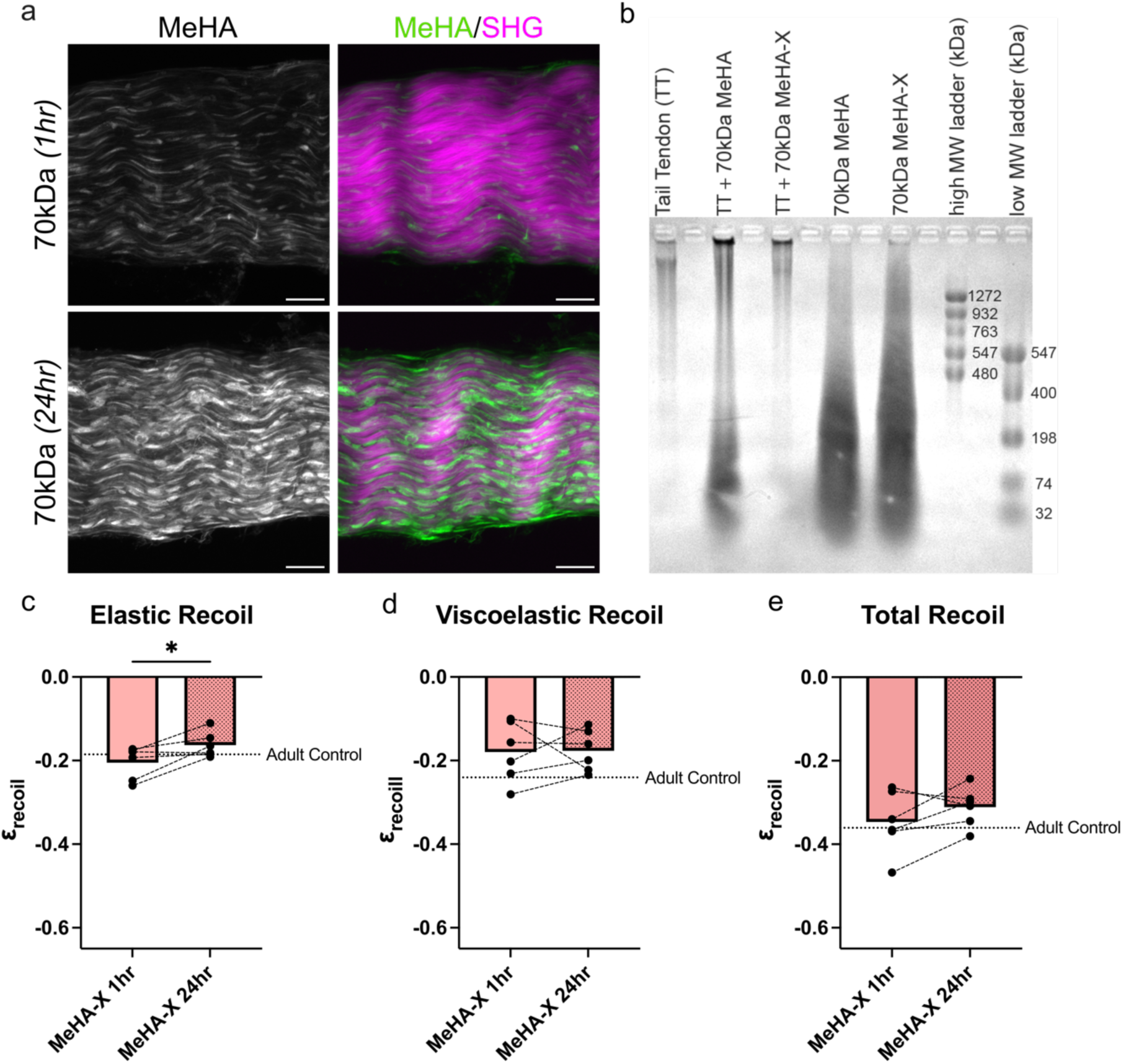
MeHA incubation time significantly affects collagen fiber elastic recoil. a) Images of tendon incubated in MeHA for 1 h and 24 h. b-d) Elastic, viscoelastic, and total recoil of ablated collagen fibers. Data represent mean with dashed lines connecting contralateral tendons from a single biological replicate. *Paired t-test*, *n =* 6 per group. e) Gel electrophoresis gel of HA in control and MeHA-treated tendons. Representative of *n* = 3 biological replicates. *p<0.05, **p<0.01, ***p<0.001.

Gel electrophoresis of HA in murine tail tendons was employed to further understand the HA network formed from crosslinking (**Figure 8E**). The MW of native HA in tendons was generally greater than 1272 kDa, which has not previously been reported and can provide guidance for HA therapeutics. Tendons in which MeHA was not crosslinked indicated that substantial MeHA was retained in the samples. With 3 minutes exposure to 455 nm light, the majority of the MeHA appeared to be crosslinked and potentially contained within the dark band observed immediately below the loading well. In addition, MeHA was crosslinked in solution using the same parameters. An overall shift in the MW of the band occurred; however, a substantial amount of MeHA remained uncrosslinked. These results further confirm that not only is the MeHA diffusing within the tendons, but the MW increases with crosslinking.

## 3. Conclusion

Our study demonstrated that HA content significantly impacts tendon collagen fiber micromechanical properties. We revealed a role of HA in tendon mechanics that has not been considered previously.^[35]^ The ability to crosslink MeHA after it has penetrated a tissue offers the ability to distribute HA into a tissue with enhanced tissue residence time, providing insight into improved delivery of HA therapeutics. MeHA treatment was able to restore the mechanical properties of aged and HAase treated tendons to that of adult control values, indicating potential mechanical benefits from using HA therapeutics in tendon. Importantly, we were able to resolve fiber-scale changes in mechanics without discernable macroscopic changes in tissue material properties. We attribute this to the development of a quantifiable measure of fiber recoil and sliding in situ via femtosecond laser ablation. Although developed for tendon, this technique is broadly applicable for investigating the micromechanical response of other collagen-based tissues, other fibrillar ECM such as elastic fibers, and engineered fibrillar hydrogels. In addition to quantifying fiber-scale mechanics, it also provides a controlled model of microscale matrix damage, enabling future investigations into of localized cellular responses.

Our method includes key improvements on capturing physiological tendon behavior. Investigating collagen fibers within the native conditions during testing, both by keeping bone and muscle insertion sites intact and by probing fibers beneath the tendon surface, enables the quantification of the impact of amorphous ECM components and surrounding fibers on the mechanical behavior collagen fibers in situ.

## 4. Experimental Section

### Materials

All reagents were purchased from Sigma Aldrich and used as received, unless otherwise noted.

### Tissue Collection

All murine experiments were approved by the University of Colorado Boulder Institutional Animal Care and Use Committee (IACUC, protocol #2705). Male and female adult (3–5-month-old) mice and aged (>16 month) mice were obtained from the Jackson Laboratory (C57BL/6J) and were phenotypically wild type. After euthanasia, the extensor carpi radialis longus (ECRL) tendon was freshly dissected from each forelimb with muscle and bone insertions attached while hydrated in 1× phosphate buffered saline (PBS). All tendons were mechanically tested the day of harvest. Tendons were then processed as described for the following studies.

### Synthesis of Methacrylated HA

Methacrylated HA was synthesized as previously described.^[83]^ Briefly, sodium hyaluronate (70 kDa; Lifecore Biomedical) was added to a round-bottom flask with a stir bar and dissolved in ice-cold deionized water. Under ice-cold conditions, NaOH was added to adjust the pH to 8.5. Methacrylic anhydride was added to the solution dropwise, alternating with dropwise addition of NaOH to maintain a pH of 8.5. Once all methacrylic anhydride was added, the pH of the solution was maintained at 7.5 to 8.5 for 4 hours through regular addition of NaOH. Solution was then stirred overnight at 4°C. The following morning, the pH was brought back to 8.5. Modified polymer was dialyzed against deionized water for 2 weeks, frozen at -80°C, and lyophilized. % modification was measured via NMR. MeHA was then functionalized with a fluorescent label for visualization. For this, MeHA was dissolved in 0.2 M triethanolamine buffer at pH 8. Thiolated FITC peptide (GCDDD-FITC; Genscript) solution (5 mM) was added dropwise to label 1% of HA repeats. Solution was mixed overnight at 37°C in a shaking incubator, dialyzed for 3 days, and lyophilized.

### Preparation of Tendons for Mechanical Testing

After tissue collection, samples underwent a three-step treatment process (**Table 1**). First, samples were incubated for 1 h at 37 °C in either 20 mM sodium acetate, pH 6 (control buffer) or the buffer with 10 U hyaluronidase (HAase) (MP Biomedicals, Hyaluronidase from *Streptomyces hyalurolyticus*) to digest endogenous HA in the tissue. Second, samples were incubated in 0.5% MeHA (70 kDa, 6% methacrylation) +/-0.05 wt/vol% lithium phenyl-2,4,6-trimethylbenzoylphosphinate (LAP), a water soluble and biocompatible photoinitiator.^[84]^ Sample groups with crosslinking were then exposed to 450 nm laser (10 mW cm^-2^) for 3 minutes (Opto Engine MDL-III-450-500mW), which can penetrate and promote crosslinking at least 250 μm into cartilage.^[75]^ Contralateral tendons from each mouse received different treatments to be compared via pairwise analysis. Preliminary experiments indicated that LAP and UV-treatment alone did not result in any significant differences in tendon mechanical properties (**Figure S2**), so both these and PBS controls were combined in control groups.

To test the effect of diffusion time on the resultant mechanical properties, we also mechanically tested contralateral tendons where one tendon was incubated in 0.5% MeHA for 1 hour and the other for 24 hours. Each group received UV-treatment and was mechanically tested immediately after incubation.

### Mechanical Testing

Mechanical testing was performed using a FUTEK LSB200 10 g high precision load cell, ThorLabs PT3 stage, and ThorLabs ZST225B actuator attached to our lab’s custom designed floating-platform system.^[50]^ Tendons were mechanically tested while imaged with an Olympus FVMPE-RS twin laser multiphoton system and a 25× (NA = 1.05) FV30-AC25W truResolution water-immersible objective using the method described in Section 2.1. Briefly, each tendon was gripped at muscle and bone insertion sites, while submersed in PBS, and displaced at a strain rate of 1% s^-1^ to a set force of 80 mN, where macroscale stress relaxation was recorded for 300 s. After, a z-stack was taken via SHG to calculate CSA. Three fiber segments (20 μm length) within the middle region of a single tendon (*i.e.* technical replicates) were ablated. The fiber recoil from each ablation was imaged for 300 s. Image sequences were used to calculate elastic, viscoelastic, and total recoil strain as well as fiber segment length used to fit a QLV model to the recoil data (see below). Fiber recoil from the three ablation technical replicates was averaged and reported for each biological replicate. Force-time data recorded from the load cell was used to calculate macroscale stress relaxation and used for subsequent QLV modeling.

### Mechanical Analysis

For fiber scale mechanical characterization, SHG image sequences were processed using FIJI to calculate average fiber segment length immediately after ablation (L_1_) and after 300 s (L_2_) to compare with the initial length (L_0_) (**Figure 1C**). These values were used to calculate the normalized change in length after load is removed, or recoil strain. The instantaneous elastic recoil strain was measured immediately after load was removed via ablation (**Equation 1**), and time-dependent viscoelastic recoil strain was measured at the end of the 300 s fiber relaxation (**Equation 2**). Total recoil strain was used to denote the total amount of fiber relaxation after ablation (**Equation 3**).

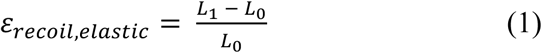

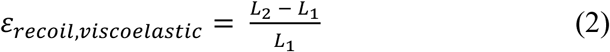

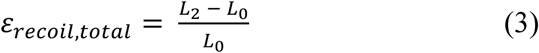

For macroscale mechanical analysis, an orthogonal section of the z-stack was used to calculate CSA by fitting an ellipse or polygon to the cross-section in FIJI. Force-displacement was processed in MATLAB to calculate the actuator displacement to the set force, defined as the distance the actuator moved from a 1 mN preload to the 80 mN set force. Force-time data was used to calculate the percent relaxation of the tissue by dividing the force at 300 s (equilibrium force) by the initial force. The relaxed stress after 300 s was calculated to compare tissue stress during ablation.

### QLV Model Fitting

A QLV model was fit to both the macroscale stress relaxation and the microscale fiber recoil after ablation. **Equation 4** shows the QLV formula, where *σ^e^*(*ε*) represents the instantaneous elastic response of the material and *G*(*t*) represents the viscoelastic relaxation function.

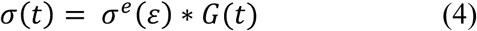

Parameters *A* and *B* are material constants in the elastic response (**Equation 5**).

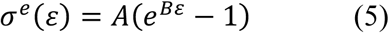

At the macroscale, a two-term relaxation function was chosen to best fit the data set, indicating two mechanisms of relaxation occurring at the tissue scale (**Equation 6**).

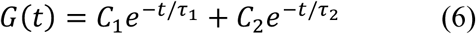

Parameters *C_1_* and *C_2_* designate relaxation ratios while τ_1_ and τ_2_ denote relaxation time constants. Fiber recoil relaxation was fit with a single-term relaxation function (**Equation 7**).

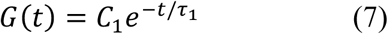

Notably, a two-term relaxation function was used for stress relaxation because a single term relaxation function did not capture the initial stress decay of the relaxation data well. In contrast, fiber recoil relaxation fitting was limited by the resolution of fiber segment length measurements in the recoil images, which dictated the number of frames sampled to eight. Therefore, a singe term relaxation function was used to avoid over-fitting the data.

### Microscopy Imaging and Analysis

To assess the role of MW on HA penetration into tendon, tendons were incubated in 0.5 wt% HA (50 kDa, 250 kDa, and 1000 kDa, Creative PEGWorks) or 70 kDa MeHA with 20% methacrylate modification for 1 hour at room temperature. Both HA and MeHA were labelled with 1% fluorescein. Tendons were immediately rinsed in PBS, fixed in 4% paraformaldehyde for 1 h, then rinsed in PBS again. Wholemount tissues were counterstained with DAPI for 48 h (1:500 dilution of 1 mg mL^-1^ stock in 0.02% bovine serum albumin in PBS). Cryosections were counterstained with DAPI for 1 h at the same dilution. All were imaged on a STELLARIS confocal microscope using the same settings across treatments at 25× with maximum z-projection shown.

Persistence of MeHA in tendon was evaluated by incubating tendons in fluorescently labeled 50 kDa HA or 70 kDa MeHA for 1 h at room temperature. HA and MeHA tendons were imaged at 0, 6, and 24 h to measure the persistence in hydrated tissue. Between timepoints, samples were stored in PBS at 4°C. Samples were wholemount imaged via confocal microscopy at 25× with maximum z-projection shown.

The effect of crosslinking was investigated by incubating tendons in 0.5% MeHA +/- 0.05 wt/vol% LAP. Samples were either kept as un-crosslinked controls or were photocrosslinked with 3 minutes laser exposure of 10 mW cm^-2^ 450 nm laser light. Tendons were imaged immediately at 25× with maximum z-projection shown or after storing in PBS at 4°C for 1 week, with daily changes of fresh PBS solution.

Endogenous HA was labeled with biotinylated hyaluronic acid binding protein (HABP) for 48 h (1:300 dilution of 1 mg mL^-1^ stock in 0.02% bovine serum albumin in PBS) then stained with Alexa Fluor 647 streptavidin for 48 h (1:500 dilution of 2 mg mL^-1^ stock in 0.02% bovine serum albumin in PBS). Collagen fibers were visualized with SHG. Samples were imaged at 25× with maximum z-projection shown.

### HA and MeHA Quantification

After mechanical testing, all samples were snap frozen in liquid nitrogen and stored at -80 °C for subsequent measurement of endogenous HA and additional MeHA content amount using an HA ELISA (Echelon Biosciences, K-1200 kit). Upon thawing, samples were dried, weighed, and incubated overnight in papain at 50 °C (60 U papain in 20 mM EDTA, 5 mM L-cysteine buffer). The enzyme was inactivated by heating samples to 100 °C for 20 minutes. Digested samples were then processed following manufacturer’s instructions, and HA amount per tissue dry weight was reported.

### HA Molecular Weight Gel Electrophoresis

Electrophoresis of HA and MeHA on agarose gel was run in 1X TBE buffer per HA ladder manufacturer instructions (Select-HA Ladders, Echelon Biosciences). Murine tail tendons were digested in digestion solution (0.15M Tris, 0.15M NaCl, 0.01M CaCl_2_, 5mM deferoxamine mesylate, and 40 units of Proteinase K) at 55 °C overnight. Before digestion, MeHA treated samples were incubated in 15% modification 70kDa MeHA for 1 hour then crosslinked if necessary. Samples were mixed with loading buffer at a 1:4 buffer to sample ratio and loaded into a 1% agarose gel. HA-Ladders were run alongside samples. Before samples were added, the agarose gel underwent pre-electrophoresis at 40V for 20 minutes. After samples were loaded, the gel was run at 20V for 30 minutes and 40V for 6 hours. The gel was stained with 0.005% (w/v) Stains-All in 50% ethanol overnight. Following staining the gel was detained using 70% ethanol overnight and imaged.

### Statistics

All statistical tests were run using GraphPad Prism 10. Brown-Forsythe one-way ANOVA was used to test for statistical significance of CSA, actuator displacement to set force, percent relaxation, and relaxed stress at 300 s. Brown-Forsythe one-way ANOVA was also used to test for statistical significance of elastic, viscoelastic, and total recoil, and uncorrected Fisher’s LSD post-hoc testing was used for pairwise comparison of groups that used contralateral limbs from a single biological replicate for separate treatments. Paired t-tests were used when only two groups from contralateral limbs were being compared, and Welch’s t-test was used to compare adult and aged tendon mechanics. Statistical significance was reported for p<0.05.

The authors acknowledge James Orth and the CU Boulder Light Microscopy Core Facility for assistance with and access to the Olympus FVMPE-RS twin laser 2-photon.

The data that support the findings of this study are available from the corresponding author upon reasonable request.

Received: ((will be filled in by the editorial staff))

Revised: ((will be filled in by the editorial staff))

Published online: ((will be filled in by the editorial staff))

## Supporting Information

Supporting Information is available from the Wiley Online Library or from the author.

## Notes

### Competing Interest Statement

The authors have declared no competing interest.

