## supporting information for "Quantifying the role of hyaluronic acid on collagen fiber mechanics via femtosecond laser ablation"

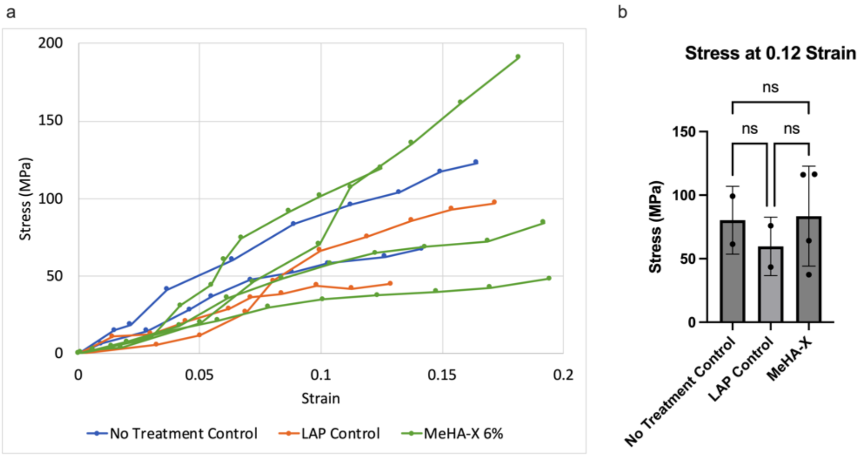


**Figure S1.** MeHA used in study (6% methacrylation) did not appear to affect tendon macroscale stiffness. a) Stress–strain curves for no treatment control, LAP control, and MeHA-X treated tendons. b) Stress at 0.12 strain. Data represent mean ± SD. Brown–Forsythe one-way ANOVA with Dunnett's T3 post hoc test, n = 2-4 per group, ns = not significant.

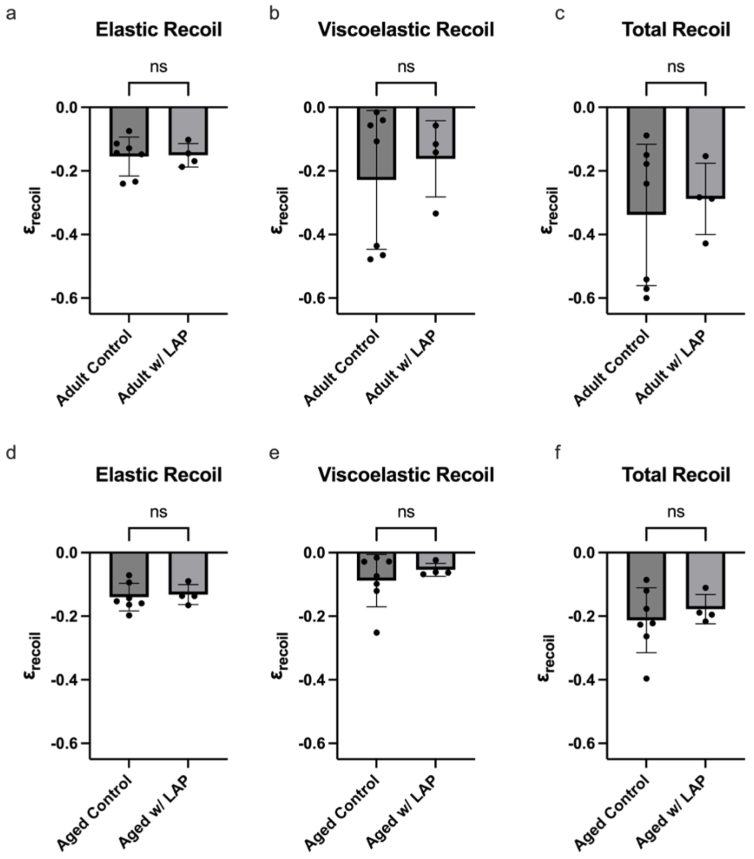


**Figure S2**. LAP and photocrosslinking did not alter collagen fiber recoil.
*a–c)* Elastic, viscoelastic, and total collagen fiber recoil following femtosecond laser ablation in adult control tendons and tendons exposed to 0.05 wt/vol% LAP with 3 min of 450 nm light exposure (Adult + LAP). *d–f)* Elastic, viscoelastic, and total collagen fiber recoil following femtosecond laser ablation in aged control tendons and tendons exposed to 0.05 wt/vol% LAP with 3 min of 450 nm light exposure (Aged + LAP). No significant differences were detected between groups (Welch's t-test, $p > 0.05$).

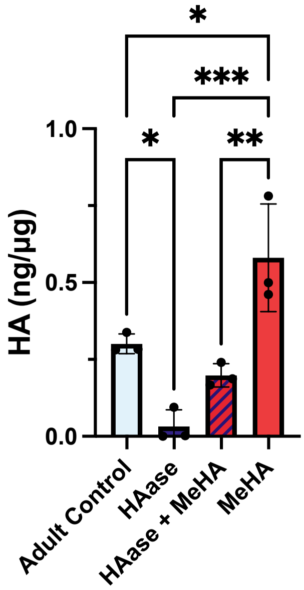


**Figure S3**: HA content per tendon dry weight measured by ELISA. HAase treatment significantly reduced tendon HA content, while MeHA supplementation restored HA levels. Data are presented as mean ± SD. HA content with treatment was significant (***p<0.001, one-way ANOVA with Dunnett's T3 multiple-comparisons test).

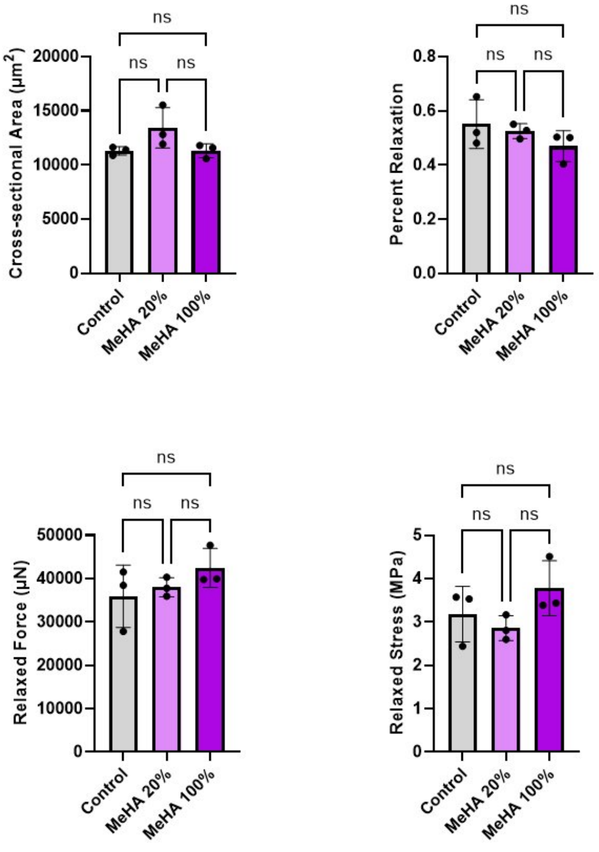


**Figure S4.** Degree of MeHA methacrylation did not significantly alter macroscale mechanical properties of tendon. Cross-sectional area, percent stress relaxation after 300 s, relaxed force after 300 s, and relaxed stress at the time of ablation. Data represent mean ± SD. Brown–Forsythe one-way ANOVA with Dunnett's T3 post hoc test, n = 3 per group. ns = not significant.

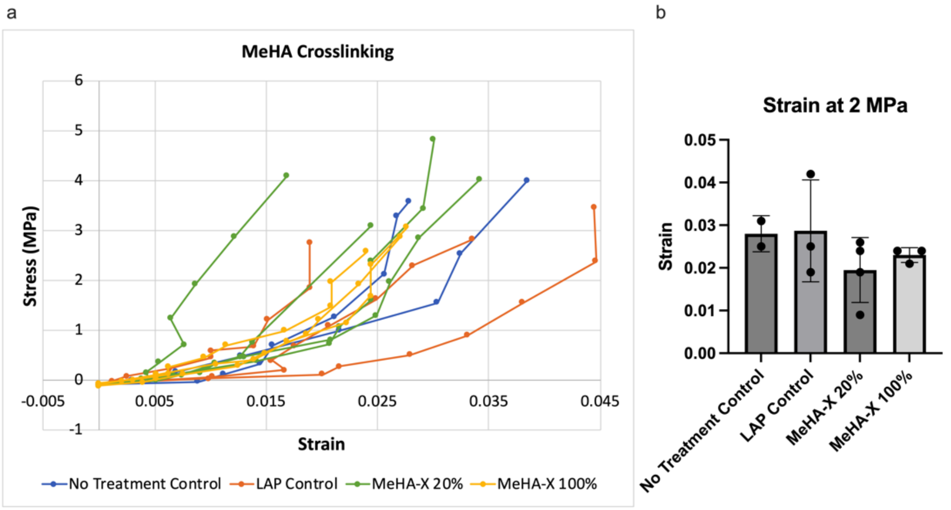


**Figure S5.** Degree of MeHA methacrylation did not significantly alter tendon tensile behavior. a) Stress–strain curves for control tendons and tendons treated with MeHA-X at varying degrees of methacrylation. b) Strain at 2 MPa stress. Data represent mean ± SD. Brown–Forsythe one-way ANOVA with Dunnett's T3 post hoc test, n = 2-3 per group. No significant differences were detected between groups (ns).

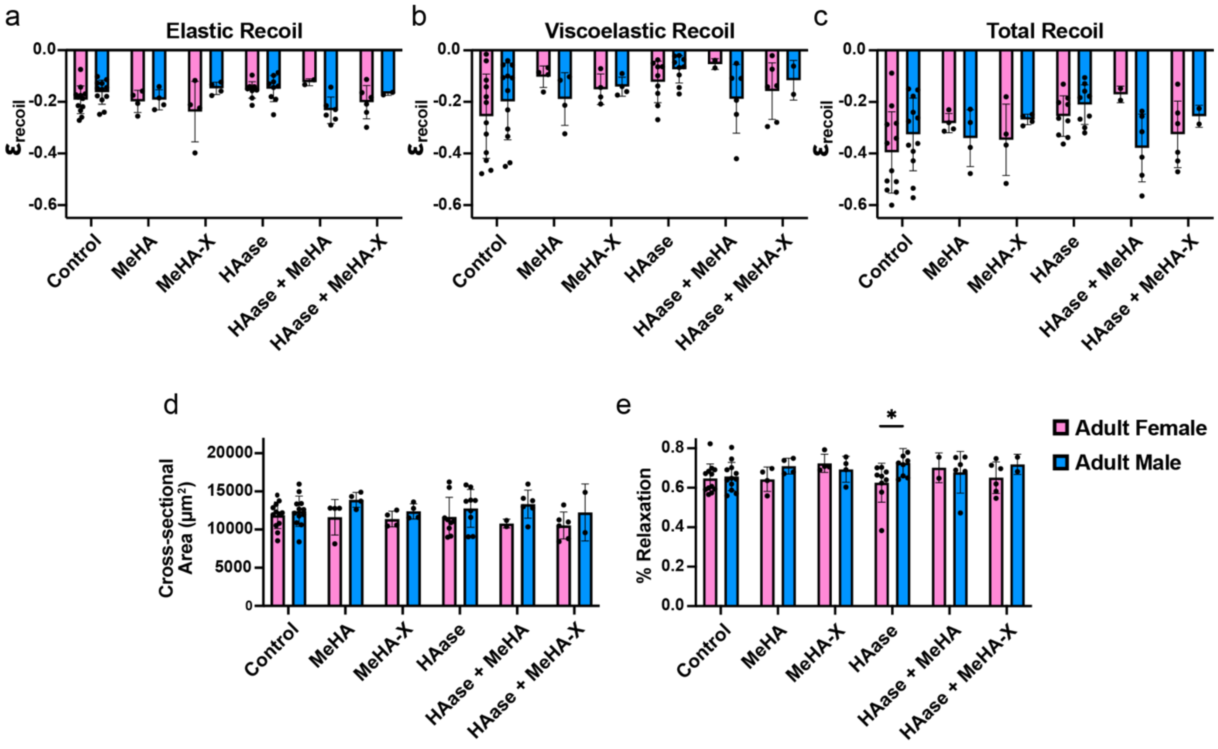


**Figure S6.** MeHA treatment altered tendon recoil behavior, while tendon cross-sectional area differed by sex. a) Elastic recoil, b) viscoelastic recoil, c) total recoil, d) cross-sectional area, and e) percent stress relaxation for adult female and adult male tendons across treatment groups. Data represent mean ± SD with individual biological replicates shown. Two-way ANOVA revealed a significant sex × treatment interaction for elastic recoil (*$p < 0.05$), significant treatment effects for viscoelastic recoil (*$p < 0.05$) and total recoil (*$p < 0.05$), and a significant effect of sex on cross-sectional area (**$p = 0.007$). n = 6–8 per group.
